# A Fast Approximate Algorithm for Mapping Long Reads to Large Reference Databases

**DOI:** 10.1101/103812

**Authors:** Chirag Jain, Alexander Dilthey, Sergey Koren, Srinivas Aluru, Adam M. Phillippy

## Abstract

Emerging single-molecule sequencing technologies from Pacific Biosciences and Oxford Nanopore have revived interest in long read mapping algorithms. Alignment-based seed-and-extend methods demonstrate good accuracy, but face limited scalability, while faster alignment-free methods typically trade decreased precision for efficiency. In this paper, we combine a fast approximate read mapping algorithm based on minimizers with a novel MinHash identity estimation technique to achieve both scalability and precision. In contrast to prior methods, we develop a mathematical framework that defines the types of mapping targets we uncover, establish probabilistic estimates of p-value and sensitivity, and demonstrate tolerance for alignment error rates up to 20%. With this framework, our algorithm automatically adapts to different minimum length and identity requirements and provides both positional and identity estimates for each mapping reported. For mapping human PacBio reads to the hg38 reference, our method is 290x faster than BWA-MEM with a lower memory footprint and recall rate of 96%. We further demonstrate the scalability of our method by mapping noisy PacBio reads (each ≥ 5 kbp in length) to the complete NCBI RefSeq database containing 838 Gbp of sequence and > 60, 000 genomes.

## 1 Introduction

Mapping reads generated by high-throughput DNA sequencers to reference genomes is a fundamental and widely studied problem [16, 24]. The problem is particularly well studied for short read sequences, for which effective mapping algorithms and widely used software such as BWA [15] and Bowtie [12] have been developed. The increasing popularity of single-molecule sequencers from Pacific Biosciences and Oxford Nanopore, and their continually improving read lengths (10 kbp and up), is generating renewed interest in long read mapping algorithms. However, the benefit of long read lengths is currently accompanied by much higher error rates (up to 15–20%). Despite their high error rates, long reads have proved advantageous in many applications including *de novo* genome assembly [7, 10] and real time pathogen identification [2, 22].

Sequence data from nanopore devices is available just minutes after introducing the sample. This can enable real-time genomic analysis when coupled with fast computational methods that can map the data stream against large reference databases. However, mapping raw sequences continues to be a bottleneck for many applications. The problem is only going to worsen as Oxford Nanopore’s PromethION is projected to generate multiple tera-bases of sequence per day. In parallel, reference databases are continually growing in size, with the non-redundant NCBI RefSeq database close to exceeding a tera-base in size. The high error rate of raw single-molecule sequences further adds to the computational complexity.

Read mapping problems can be solved exactly by designing appropriate variants of the Smith-Waterman alignment algorithm [27]; however, it is computationally prohibitive when mapping reads from a high throughput sequencer to large reference genomes. Seed-and-extend mapping heuristics address this limitation for both long and short reads by limiting the search to locations where exact short word or maximal common substring matches occur before executing an alignment algorithm at these locations [1, 8, 5]. Accurate alignment-based long read mappers include BLASR [5] and BWA-MEM [13]. However, repetitive seeds that do not translate to correct mappings combined with high sequencing error rates limit their scalability. Additionally, alignment-based mapping algorithms preserve the complete reference sequence in the index, and hence, cannot scale to tera-base scale reference databases. Many genomics applications do not require detailed base-to-base alignment information, and instead use only the alignment boundary and identity summaries. Such applications include depth-of-coverage analysis, metagenomic read assignment, structural variant detection, and selective sequencing [18]. Efficient algorithms for these problems, combined with nanopore sequencing, could enable the real-time genomic analysis of patients, pathogens, cancers, and microbiomes.

One class of algorithms for fast, approximate mapping relies on ideas originally developed for finding similarities between web documents. Broder [4] proved that an unbiased estimate of the Jaccard similarity coefficient between two sets can be computed efficiently using a subset of hashed elements called a sketch. Schleimer *et al*. [25] proposed the winnowing algorithm, which picks a minimum hashed item (also known as a minimizer [23]) from each consecutive window of text as a means to more quickly estimate local similarity between web documents. These ideas have been used to develop new mapping and assembly algorithms for long reads such as the MinHash Alignment Process [3], minimap [14], and BALAUR [21]. To date, the effectiveness of these approaches has only been demonstrated empirically.

In this paper, we propose a fast approximate algorithm for mapping long reads that scales to large reference databases with sufficient theoretical guarantees and practical validation on the quality of results reported. We propose a problem formulation that mathematically characterizes desired mapping targets by linking the Jaccard coefficient between the *k*-mer spectra of the read and its mapping region to a sequence error rate assuming a Poisson error model. We then provide an efficient algorithm to estimate the Jaccard coefficient through a combination of MinHash and winnowing techniques that characterizes and guarantees the types of mapping regions we find. On the quality side, we provide probabilistic bounds on sensitivity. We present techniques for choosing algorithmic parameters as a function of error rate and sequence lengths that guarantees the desired statistical significance. The theory is validated using PacBio and MinION datasets, and we demonstrate the scalability of our approach by mapping PacBio metagenomic reads to the entire RefSeq database. The speed and space efficiency of our algorithm enables real-time mapping, and compared to minimap, our method maintains high sensitivity with better precision for large, repetitive genomes. The implementation is available at github.com/MarBL/MashMap.

## 2 Preliminaries

### Read Error Model

We assume errors occur independently at the read positions, and use a Poisson error model as in previous works [9, 19]. A binomial model would also be appropriate, but is not discussed here for brevity. Let ϵ ∈[0, 1] be the per-base error rate. The expected number of errors in a *k*-mer is *k · ϵ*, and the probability of no errors within each *k*-mer, assumed independent, is *e*^−^*^ϵk^*. We assume the statement is valid irrespective of error type.

### Jaccard Similarity

Assuming 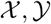 are the sets of *k*-mers in sequences *X* and *Y* respectively, their Jaccard similarity is defined as 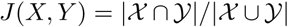. The Poisson error model is used to compute the relationship between Jaccard similarity and alignment error rate [19]. We approximate the length of a read alignment to be the read length. Let *A* be a read derived from *B_i_*, where *B_i_* denotes the length |*A*| substring of reference *B* starting at position *i*. If *c* and *n* denote the number of error-free and total *k*-mers in *A*, respectively, then the expected value of *c/n*, termed *k-mer survival probability*, is *e*^-^*^ϵk^*. This equation assumes *k* is large enough such that *k*-mers in *A* or *B_i_* are unique. Because |*A*| = |*B_i_*|, *J* (*A, B_i_*), abbreviated as *J*, equals *c*/ (*2n*-*c*). Using the two equations, we derive the following functions 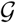 and 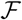 to estimate *J* and *ϵ*:

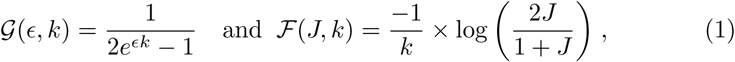
where 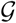(*ϵ, k*) serves as an estimate of the Jaccard similarity given an error rate and 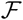(*J, k*) estimates the converse. Note 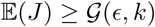 (using Jensen’s inequality).

### MinHash Approximation

The MinHash algorithm is a fast and space-efficient approximation technique to compute an unbiased estimate of Jaccard similarity [4], without explicitly computing the underlying set intersection and union. Let *s* be a fixed parameter. Assuming universe *U* is the totally ordered set of all possible items, and *Ω*: *U* → *U* is a permutation chosen uniformly at random, Broder [4] proved that 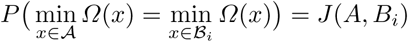, and that
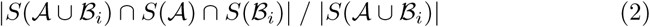
is an unbiased estimate of *J*(*A, B_i_*), where *S*(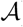) (called the *sketch* of 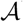) is the set of the smallest *s* hashed items in 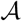, i.e., *S*(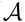) = min*_s_*{Ω(*x*): *x* ϵ 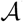}. Typically, the denominator 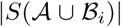 is referred as the MinHash sketch size and the numerator as the count of shared sketch elements. This estimate is unbiased provided *S*(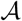) is a simple random sample of 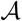. Increasing the sketch size improves the accuracy of the estimate.

Assuming *s* is fixed and the true Jaccard similarity *j* = *J* (*A, B_i_*) is known, the count of shared sketch elements between *S*(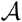) and *S*(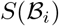) follows a hypergeometric distribution. Since *s* is much smaller than 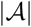, it can be approximated by the binomial distribution.

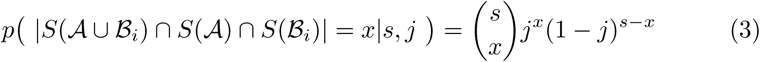
As an example, Figure 1 illustrates this distribution for a read with known Jaccard similarity *j* = 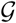(ϵ = 0.15, *k* = 16) (using Equation 1) and sketch size *s* varying from 200 to 500.

**Fig.1.**
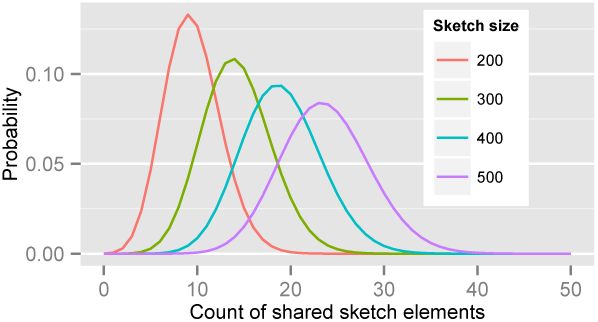
Probability distributions of count of shared sketch elements for a read with 15% alignment error (*ϵ* = 0.15) and *k*-mer size of 16, with varying sketch sizes. Estimated Jaccard similarity computed using Equation 1 is 0.0475.

### Winnowing

Winnowing is a local fingerprinting algorithm, proposed to measure similarity between documents by using a subset of hashed words [25]. Unlike MinHash sketching, it bounds the maximum positional gap between any two consecutive selected hashes. It works by sampling the smallest hashed item in every consecutive fixed size sliding window (Figure 2). Formal description of this algorithm in the context of genomic sequences follows.

**Fig.2.**
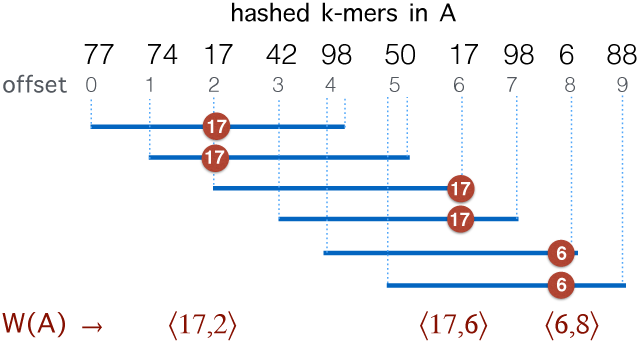
Illustration of the winnowing method on a sequence of hashed *k*-mers in *A. W* (*A*) represents the minimizers sampled from the sequence with window size *w* = 5.

Let *A*_0_ denote the set of all *k*-mer tuples 〈*k_i_, i*〉 in sequence *A, i* denoting the *k*-mer position. Let *w* be the window-size used for winnowing, and *K_j_* be the set of *w* consecutive *k*-mer tuples starting at position *j* in *A*, i.e., *K_j_* = {〈*k_i_, i*〉: *j* ≤ *i* < *j* + *w*}. Assume *Ω* is a hash function defined as a random permutation. Then, the set of *minimizers* sampled by the winnowing algorithm in sequence *A* is 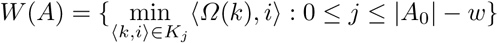 where

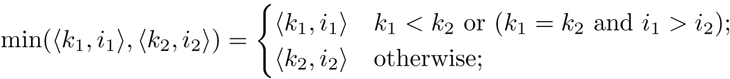
Schleimer *et. al*. [25] prove that the expected set count of minimizers selected from a random sequence *A* is 2|*A*_0_|/*w*. Moreover, *W*(*A*) can be computed efficiently in *O*(|*A*|) time and *O*(*w*) space using a double-ended queue, as sequence *A* is read in a streaming fashion [26].

### 3 Problem Formulation

Given a read *A* and the maximum per base error rate *ϵ_max_*, our goal is to identify target positions in reference *B* where *A* aligns with ≤ *ϵ_max_* per-base error rate. This problem can be exactly solved in *O*(|*A*|·|*B*|) time by designing a suitable quadratic time alignment algorithm. When mapping to a large database of reference sequences, solving this problem exactly is computationally prohibitive. Hence, we define an approximate version of this problem using the Jaccard coefficient as a proxy for the alignment as follows: Let *B_i_* denote the substring of size |*A*| in *B* starting at position *i* (0 ≤ *i* ≤ |*B*| – |*A*|). For a given *k*, we seek all mapping positions *i* in *B* such that

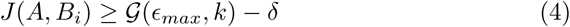
Note that if *A* aligns with *B_i_* with per-base error rate *≤ ϵ_max_*, then 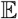( *J* (*A, B_i_*)) ≥ 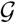(ϵ*_max_*,*k*) (using Equation 1). As this equation applies only to the expected value of *J* (*A, B_i_*), we lower this threshold by *δ* to account for variation in the estimate. The parameter *δ* is defined as the margin of error in Jaccard estimation using a 90% confidence interval.

## 4. The Proposed Algorithm

Directly computing *J*(*A, B_i_*) for all positions *i* is as asymptotically expensive as the alignment algorithm. The rationale for reformulating the problem in terms of Jaccard coefficients is that it enables the design of fast heuristic algorithms. We present an algorithm to estimate *J*(*A, B_i_*) efficiently using a combination of MinHash and winnowing techniques. In addition, we compute an estimate of the alignment error rate *ϵ* for each mapping reported. Our method relies on an indexing and search strategy we developed to prune the incorrect mapping positions efficiently.

### 4.1 Definitions

Let *W*(*A*) be the set of minimizers computed for read *A* using the winnowing method with window-size *w*. We sketch *W*(*A*) instead of sketching *A* itself. Assuming s is a fixed parameter, we define *S*(*W*(*A*)) as the set of the *s* smallest hashed *k*-mers that were sampled using winnowing of *A*, i.e., *S*(*W*(*A*)) = min*_s_*{*h*: 〈*h, pos*〉 ∈ *W* (*A*)}. To estimate *J*(*A, B_i_*), we define *winnowed-minhash* estimate 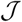 (*A, B_i_*) for *J* (*A, B_i_*) as

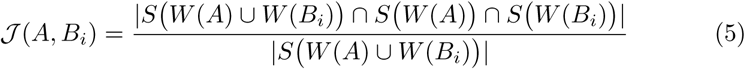
In contrast to the MinHash approximation (Equation 2), our estimator 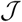 (*A, B_i_*) uses winnowing to reduce the sampling frame before picking the minimum hash values. Even though *S*(*W*(*A*)) is no longer a simple random sample of the *k*-mers in *A*, we empirically show in Section 8.1 that the quality of the Jaccard estimation using 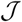(*A, B_i_*) is as good as the MinHash estimation. We use *W_h_*(*A*) to denote the set of hashed *k*-mers in *W*(*A*), i.e., *W_h_*(*A*) = {*h*: 〈*h, pos*〉 ∈ *W*(*A*)}.

### 4.2 Indexing the Reference

Retaining the minimizers *W*(*B_i_*) is sufficient for Jaccard similarity estimation 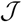 (*A, B_i_*) (Equation 5). Since *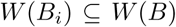* (Section 2), we compute *W* (*B*) from the reference sequence *B* in order to be able to extract *W*(*B_i_*) efficiently for any *i*. The set *W*(*B*) can be computed from *B* in a linear scan in *O*(|*B*|) time. We store *W*(*B*) as an array 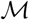 of tuples 〈*h, pos*〉. When created, the set is naturally in ascending sorted order of the positions. Further, to enable *O*(1) look-up of all the occurrences of a particular minimizer’s hashed value *h*, we also replicate *W*(*B*) as a hash table 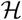 with *h* as the key and an array of its positions {pos: 〈*h*,*pos*〉 *ϵ W*(*B*)} as the mapped value. The expected space requirements for 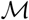 and 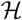 are 2|*B*|/*w* (Section 2). We postpone our discussion on how to compute an appropriate window-size *w* to Section 5. Besides low memory requirements, a key advantage of this indexing strategy is that a new reference sequence can be incrementally added to the existing data structure in time linear to its length, which is not feasible for suffix array or Burrows-Wheeler transform based indices, typically used in most mapping software.

### 4.3 Searching the Reference

The goal of the search phase is to identify for each read *A*, positions *i* such that *J* (*A, B_i_*) ≥ 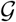(*ϵ_max_, k*) – *δ*. We instead compute the winnowed-minhash estimate 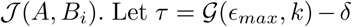. To avoid directly evaluating 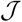 (*A, B_i_*) for each *B_i_*, we state and prove the following theorem:

#### Theorem 1.

Assuming sketch size *s* ≤ |*Wh*(*A*)|,

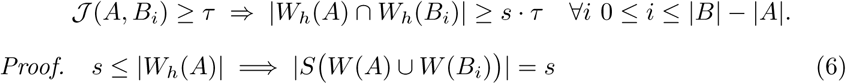
From Equation 5,

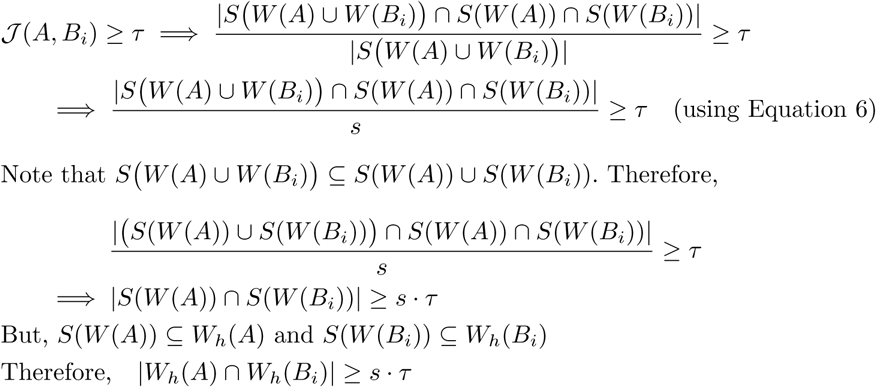

We use the above condition as a filter and only consider positions in *B* which satisfy |*W_h_*(*A*) ∩ *W_h_*(*B_i_*)| ≥*s* · *τ*.To maximize effectiveness of the filter, we set the sketch size *s* = |*W_h_*(*A*)|. The search proceeds in two successive stages. The first stage identifies candidate positions *i* using Theorem 1, and the second stage computes 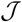(*A, B_i_*) at each candidate position *i*. The position is retained as output if 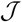(*A, B_i_*) ≥ *τ*, and discarded otherwise.

Stage 1:Algorithm 1 outlines the first stage of our mapping procedure. It calculates all offset positions *i* in *B* such that |*W_h_*(*A*)∩*W_h_*(*B_i_*)| ≥ 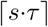 = *m*. The output list *T* is created in the form of one or more tuple ranges 〈*x, y*〉, implying that the criterion holds true for all *B_i_, x* ≤ *i* ≤ *y*. We begin by computing the minimizer hashed values *W_h_*(*A*) by winnowing the read *A*, and compute the positions of their occurrence in the reference (line 4). Accordingly, *L* = {*pos*: *h* ∈ *W_h_*(*A*) ^ 〈*h, pos*〉∈ *W*(*B*)}. Next, we sort the array *L* to process all the positions in ascending order. If *B_i_* satisfies the filtering criterion, there should be at least *m* entries in *L* with values between [*i, i* + |*A*|). It also implies that m consecutive entries should exist in *L* with positional difference between the first and *m^th^* entry being < |*A*|. This criterion is efficiently evaluated for all *B_i_* using a linear scan on *L* (lines 6–9). If satisfied, we push the associated candidate range into *T*. To avoid reporting *B_i_* more than once, we merge two consecutive overlapping tuple ranges into one.

#### Algorithm 1: Stage 1 of mapping read

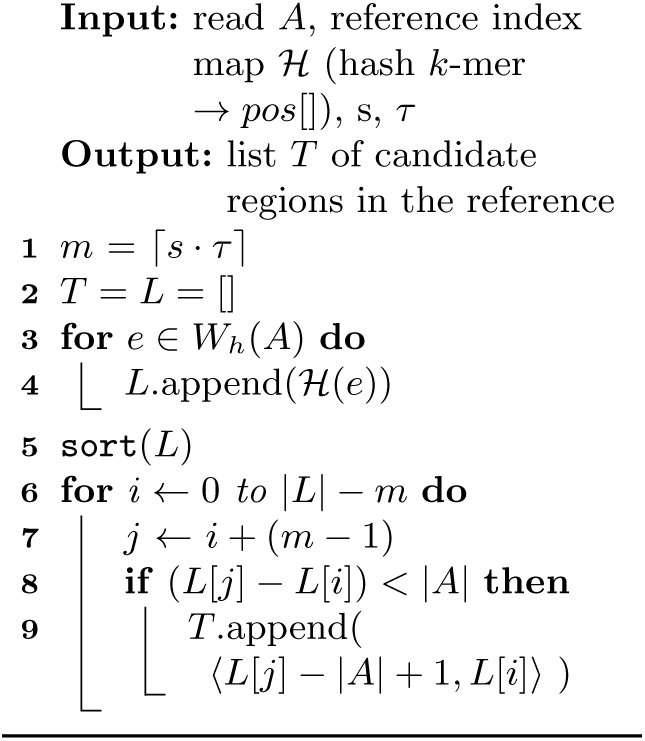

#### Algorithm 2: Stage 2 of mapping read

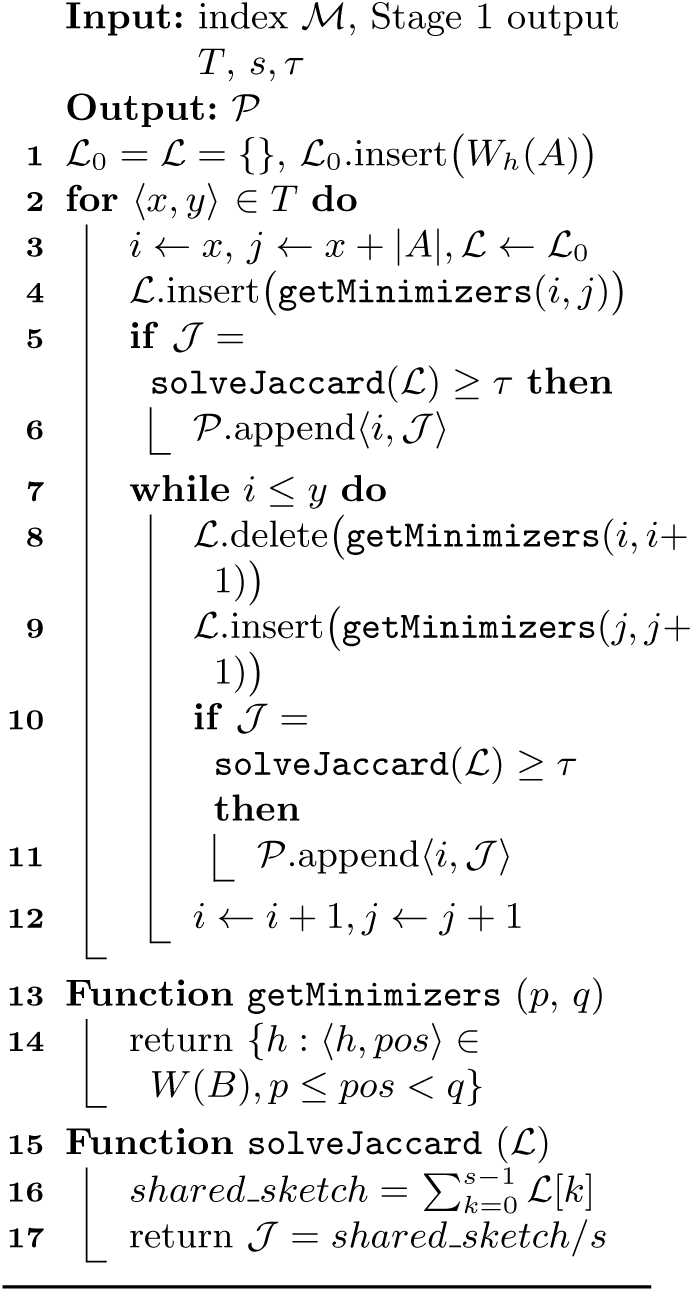

Stage 2:Evaluation of each tuple 〈*x, y*〉 in the Stage 1 output array *T* requires computing 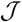 (*A, B_i_*) ∀*i, x* ≤ *i* ≤ *y*. Accordingly, we compute the minimum *s* unique sketch elements within *W_h_*(*A*) ∪ *W_h_*(*B_i_*), and count the ones shared between *A* and *B_i_*. We show the step-by-step procedure in Algorithm 2. We use 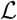 to contain the minimizer hashed values {*h ϵW_h_*(*A*) ∪ *W_h_*(*B_i_*)}. To implement 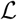, we make use of the C++ ordered map data structure that supports logarithmic time insertion, deletion and linear time iteration over unique ordered keys. We keep the hashed value as the map’s key, and map it to 1 if it appears in both the reference and the read, and 0 otherwise. For each tuple 〈*x, y*〉, we begin by saving the hashed values *W_h_*(*A*) in read *A* into map 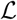 (lines 1, 3). Two loops (lines 2, 7) evaluate each tuple 〈*x, y*〉 in *T*, and consider each *B_i_, x* ≤ *i* ≤ *y* for Jaccard estimation 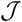 (*A, B_i_*). The function `getMinimizers` gathers the reference minimizer hashes *W_h_*(*B_i_*) by sequentially iterating over 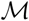 in the required position range and populating the minimizers associated with each *B_i_* into the map 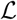 (lines 4,8-9). Note that a few incorrect corner minimizers {*h*: 〈*h, pos*〉 ∈ *W*(*B*), *i* ≤ *pos* ≤ *i* + |*A*|}\*W_h_*(*B_i_*) can appear in 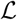 that were winnowed from windows overlapping with *B_i_*. However, these can be discarded by recomputing the minimum of the first and last window of *B_i_*. Finally, function `solveJaccard` computes |*S*(*W* (*A*)∪*W* (*B_i_*)) ∩ *S* (*W* (*A*)) ∩ *S* (*W* (*B_i_*))| by iterating over *s* minimum unique sketch elements and counting the ones shared between *A* and *B_i_*. If 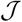(*A, B_i_*)≥ *τ*, then the position *i* and Jaccard estimate 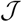 (*A, B_i_*) are saved into the output 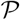 as pair 〈*i*,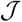(*A, B_i_*)〉The corresponding estimate of the alignment error rate *ϵ* in this case, computed using Equation 1, would be 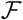(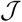(*A, B_i_*),*k*).

## 5 Selecting Window and Sketch Sizes

The sketch size for Jaccard similarity estimation is inversely proportional to the window size *w* (Section 4.3). A larger window size improves the runtime and space requirement during the search but also negatively affects the statistical significance and accuracy of our estimate. To achieve the right balance, we analyze the p-value of a mapping location being reported under the null hypothesis that both query and reference sequences are random. For the subsequent analysis, we will assume the sketch size is *s*, the count of shared sketch elements is a discrete random variable *Z*, the *k*-mer size is *k*, the alphabet set is *Σ*, and the read and reference sequence sizes are *q* and *r* respectively.

Location *i* is reported if 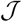(*A, B_i_*) ≥ *τ*, i.e., at least 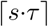 sketch elements are shared. Following [19], consider two random sequences of length *q* with *k*-mer sets *X* and *Y* respectively. The probability of a random *k*-mer *α* appearing in *X* or *Y*, assuming *q* ≫ *k*, is 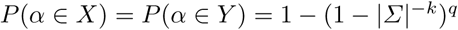. Therefore, the expected Jaccard similarity *J_null_* = *J*(*X, Y*) is given by

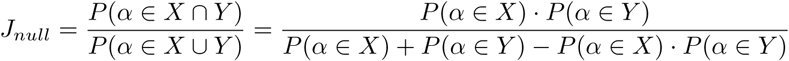
For sketch size *s*, the probability that *x* or more sketch elements are shared is 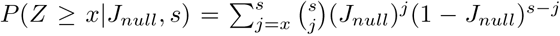. Using this equation, we compute the probability of a random sequence of length *q* mapping to at least one substring in a random reference sequence of size *r* ≫ *q* as 1 – (1 – *P*(*Z* ≥ *x*|*J_null_, s*))^*r*^. For a minimum read length *l*_0_ and 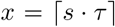 we wish to ensure that this probability is kept below a user-specified threshold *p_max_*. As reported mapping locations *i* must satisfy 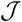 (*A, B_i_*) ≥ *τ* and *q* ≥ *l*_0_, a mapping with 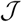 (*A, B_i_*) = *τ, q* = *l*_0_, in general, will have the highest probability of generating a random match. Therefore, we compute the maximum value of *w* that satisfies the *p_max_* constraint for this instance. Sketch size *s* is set to |*W_h_* (*A*)|, which from Section 4.3 is expected to be *q*·2/*w*. Since *x, s*, and *w* have a circular dependency, we iteratively solve for *w*, starting from the maximum value *l*_0_, until the probability of a random mapping is ≤ *p_max_*. Influence of different parameters on window size is shown in Figure 3. The window size *w* increases with increasing *p_max_* or *l*_0_, but has an inverse relationship with *ϵ_max_*. These plots also highlight that as read length and error rate improve, our algorithm automatically adapts to a larger window size, greatly improving efficiency.

**Fig. 3.**
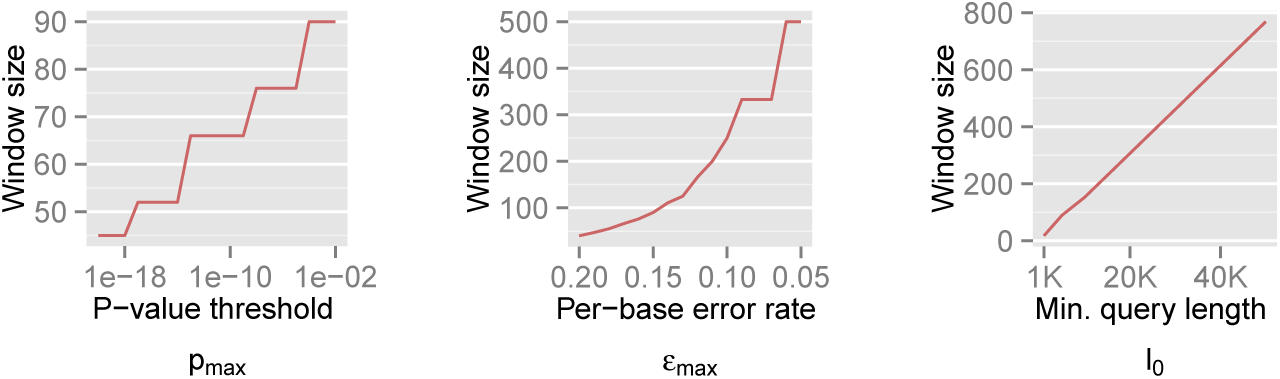
Illustration of how *w* varies with *p_max_, ϵ_max_*, and *l*_0_, respectively. The default values are set as *l_0_* = 5000, *ϵ_max_* = 0.15, *p_max_* = 0.001, *k* = 16, and *r* = 10^9^. Steps appear in the first two curves because *Z* is a discrete variable.

## 6 Proof of Sensitivity

We analyze the sensitivity exhibited by our algorithm in identifying correct mapping locations as a function of the read alignment error rate. Let *i* be a correct mapping location for read A. If *ϵ_true_* is the true error rate in aligning *A* with *B_i_*, then *J_true_* ≈ 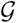(*ϵ_true_, k*). Our algorithm reports this mapping location if the Jaccard estimate 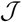 (*A, B_i_*) ≥ *τ*, i.e., the count of shared sketch elements *Z* ≥ *s* · *τ*. The associated probability is given by *P*(*Z* ≥ *s* · *τ*| *J_true_,s*) ≈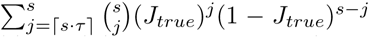. We report the corresponding values in Table 1 while varying *ϵ_max_* and *ϵ_true_* from 0.04 to 0.20 error rate, for two sketch sizes *s* = 200 and 500, respectively. In an ideal scenario, a mapping should be reported only if *ϵ_true_* ≤ *ϵ_max_*, i.e., a perfect algorithm would have “1” in each of the entries at or above the diagonal, and “0” in all other positions. From the table, it is evident our algorithm achieves close to ideal sensitivity for alignment error rates up to 20%.

**Table 1.** Probability of a mapping location being reported by our algorithm for different values of *ϵ_true_* and *ϵ_max_*. True mapping locations correspond to *ϵ_true_* ≤ *ϵ_max_*, i.e., entries at or above the diagonal in the tables. Sketch sizes are set to 200 and 500 for the left and right tables, respectively. The *k*-mer size *k* is set to 16.

| $\epsilon_{true}$ | $\epsilon_{max}$ | | | | |
| --- | --- | --- | --- | --- | --- |
|  | 0.04 | 0.08 | 0.12 | 0.16 | 0.20 |
| 0.04 | 0.951 | 1 | 1 | 1 | 1 |
| 0.08 | 0 | 0.937 | 1 | 1 | 1 |
| 0.12 | 0 | 0.016 | 0.925 | 1 | 1 |
| 0.16 | 0 | 0 | 0.184 | 0.907 | 0.997 |
| 0.20 | 0 | 0 | 0.003 | 0.403 | 0.922 |
|  | 0.04 | 0.08 | 0.12 | 0.16 | 0.20 |
| 0.04 | 0.939 | 1 | 1 | 1 | 1 |
| 0.08 | 0 | 0.949 | 1 | 1 | 1 |
| 0.12 | 0 | 0 | 0.937 | 1 | 1 |
| 0.16 | 0 | 0 | 0.013 | 0.904 | 1 |
| 0.20 | 0 | 0 | 0 | 0.104 | 0.896 |

## 7 Other Implementation Details

### Optimizing for Variable Read Lengths

In contrast to cyclic short-read sequencing, single-molecule technologies can generate highly variable read lengths (e.g. 10^2^–10^5^ bases) Previously, we discussed how the window size *w* is determined using the minimum read length *l*_0_ in Section 5. From Figure 3(c), notice that we can further reduce the sampling rate (i.e. use a larger window size) for reads longer than *l*_0_ while still satisfying the p-value constraint. However, to realize this, the sampling scheme for indexing the reference sequence B needs to be consistent with that of query. We propose the idea of *multilevel winnowing* to further optimize the runtime of our algorithm by choosing custom window size for each input read. Suppose *W_w_* (*B*) denotes the set of winnowed fingerprints in the reference computed using window size *w*, then 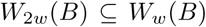 [25]. We exploit this property to construct a multilevel reference index with multiple window sizes {*w*, 2*w*, 4*w*…} recursively. This optimization yields us faster mapping time per base pair for reads longer than *l*_0_ as we independently compute the window size for a given read length *l* ≥ *l*_0_, and round it to the closest smaller reference window size {*w*,2*w*,4*w*…}. The expected time and space complexity to index the reference using multiple levels is unaffected because the expected size of 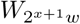(*B*) is half of 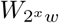 (*B*) and 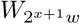 (B) can be determined in linear time from 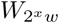 (*B*).

### Strand Prediction

To account for the reads sequenced from the reverse strand relative to the reference genome, we compute and store only canonical *k*-mers, i.e. the lexicographically smaller of the forward and reverse-complemented *k*-mer. For each *k*-mer tuple 〈*k, i*〉 in *W*(*A*) and *W*(*B*), we append a strand bit 1 if the forward k-mer is lexicographically smaller and –1 otherwise. While evaluating the read mappings in Stage 2, we compute the mapping strand of the read through a consensus vote among the shared sketches using sum of pairwise products of the strand bits.

## 8 Experimental Results

### 8.1 Quality of Jaccard Estimation

We first show that the accuracy of the *winnowed-minhash* estimator 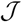 to estimate the Jaccard similarity is as good as the direct MinHash approximation, which is an unbiased statistical estimator. We construct a random sequence of length 5 kbp with each character having equal probability of being either A,C,G or T. We generate reads while introducing substitution errors at each position with probability 0.15. Note that both substitutions and indels have a similar effect of altering the *k*-mers containing them, and a uniform distribution of errors alters more *k*-mers than a clustering of errors. Figure 4 shows the estimation difference against the true Jaccard similarity using MinHash and our estimator for two different sketch sizes *s* = 100 and *s* = 200. Based on these results, we conclude that the bias in our estimation is practically negligible as the mean error by our method in estimating Jaccard similarity is < 0.003 for both sketch sizes. Similar to MinHash approximation, we note that the magnitude of estimation error reduces with increasing sketch size.

**Fig.4.**
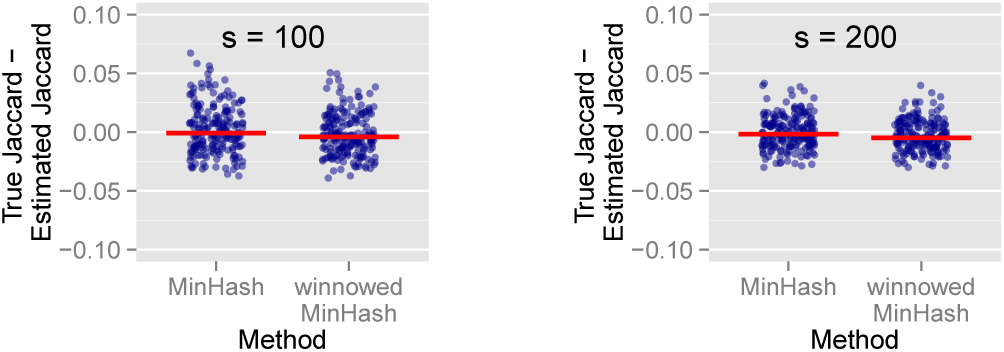
Jaccard similarity estimation using MinHash and *winnowed-minhash* estimator 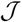 (*A, B_i_*) over simulated reads, with sketch sizes *s* = 100 and *s* = 200. Red bar indicates the average estimation difference over all reads.

### 8.2 Mapping MinION and PacBio Reads

We refer the C++ implementation of our algorithm as mashmap and compare its run-time performance and memory usage against alignment based long-read mappers BWA-MEM (v0.7.15-r114) [13], BLASR (vSMRTportal 2.3.0) [5], and minimap (v0.2) [14]. We also perform a comparison of the approximate mapping targets generated by mashmap and minimap. Like mashmap, minimap uses winnowing to index the reference, but does not use the MinHash approximation to estimate Jaccard similarity or nucleotide identity. Instead, minimap seeks clusters of minimizer matches to identify regions of local similarity. Importantly, minimap approximates a local alignment process, which is useful for split-read mapping. However, because mashmap is currently designed to find complete read mappings, we only consider this case for the following comparisons.

### Datasets and Methodology

We evaluated the algorithms by mapping long read datasets generated using single-molecule sequencers from Pacific Biosciences and Oxford Nanopore, and report single-threaded execution timings on an AMD Opteron 2376 CPU with 64 GB RAM. We use two datasets, labeled N1 and P1 respectively, both containing reads of length ≥ 5kbp. Dataset N1 is a random sample of 30,000 reads from the MinION (R9/1D) sequencing dataset of the *Escherichia coli* K12 genome [17]. Dataset P1 contains 18,000 reads generated through a single SMRT cell from PacBio’s (P6/C4) sequencing of the human genome (CHM1) [6]. We map N1 to *E. coli* K12 (4.6 Mbp) and P1 to the human reference (3.2 Gbp). For mashmap, we use the following parameters: *l*_0_ = 5000, *ϵ_max_* = 0.15, and *p_max_* = 0.001. When a read maps to multiple locations, mashmap only reports locations where mapping error rate is no more than 1% above the minimum of error rate over all such locations.

### Run-Time Performance

Run-times for the index building and mapping stages, and memory used, for the four methods are compared in Table 2. As both BWA-MEM and BLASR are alignment based methods, we expect their run-times to be significantly higher. Indeed, they take several hours in comparison to seconds (N1) or a few minutes (P1) taken by mashmap and minimap. The principal challenge is whether the latter methods can retain the quality obtainable through alignment based methods. We note that mashmap has the lowest memory footprint for both datasets, and its run-time compares favorably with minimap. The ability to compute the sampling rate at runtime gives mashmap its edge in terms of memory usage.

**Table 2.** Runtime and memory usage comparison of mashmap against minimap, BWA-MEM and BLASR for N1, P1 datasets BWA-MEM was executed with long read mapping parameters -x `pacbio/ont2d`

| Method | N1 (MinION-K12) |  |  | P1 (Pacbio-CHM1) |  |  |
| --- | --- | --- | --- | --- | --- | --- |
|  | Index | Map | Memory (MB) | Index | Map | Memory (GB) |
| mashmap | **0.5s** | 54s | **17** | 5m 52s | **1m 24s** | **3.7** |
| minimap | 0.7s | **37s** | 232 | **3m 7s** | 1m 56s | 6.8 |
| BWA-MEM | 2.6s | 5h 39m | 72 | 1h 19m | 6h 46m | 5.5 |
| BLASR | 1.3s | 10h 17m | 697 | 40m 36s | 20h 40m | 17.6 |

### Quality of Mapping

As there is no standard benchmark using real datasets, we assess sensitivity/recall using BWA-MEM’s starting read mapping positions, and precision by computing Smith-Waterman (SW) alignments of the reported mappings (Table 3). Since both minimap and BWA-MEM also report split-read alignments, we post-filter their results to only keep alignments with ≥80% read coverage. Recall is measured against BWA-MEM alignments which satisfy the *ϵ_max_* = 0.15 cutoff (≥85% identity). Because both minimap and mashmap estimate mapping positions, the reported mapping is assumed equivalent to BWA-MEM if the predicted starting position of a read is within ±50% of its length. Precision was directly validated using Smith-Waterman (SW) alignment (with scoring matrix: *match* = 1, *mismatch* = –1, *gap_open_* = –2, *gap_extend_* = –1). For minimap’s and our results, we allow SW-identity ≥ 75% and query coverage ≥ 80%. Results in Table 3 show that both mashmap and minimap have close to ideal sensitivity/recall, demonstrating their ability to uncover the right target locations.

**Table 3.** Precision and recall statistics of mashmap and minimap using datasets N1 and P1.

| Id | Recall statistics |  |  | Precision statistics |  |
| --- | --- | --- | --- | --- | --- |
|  | mashmap | minimap | #BWA mappings | mashmap | minimap |
| N1 | **100%** | 99.87% | 10,823 | **94.39%** | 94.32% |
| P1 | 96.8% | **98.7%** | 10,115 | **84.59%** | 30.34% |

Mashmap also achieves high precision, avoiding false positives on the repetitive human genome. Minimap’s low precision on human is largely driven by false-positive mappings to repetitive sequence, which could potentially be resolved with alternative clustering parameters. Mashmap false positives are dominated by reported mappings with a SW query coverage less than 80% of the read length. It may be possible to avoid such mappings by considering the positional distribution of shared sketch elements during the second stage filter, or by adopting a local alignment reporting strategy like minimap.

We compare our identity estimates (1 – *ϵ*) × 100 against the SW alignment identities in Figure 5. For the PacBio reads, we observe that most of the points are aligned close to *y* = *x*. However, for the nanopore reads, our approach overestimates the identity. This is because PacBio sequencing produces mostly random errors, whereas current nanopore errors are more clustered and systematic [11].

**Fig. 5.**
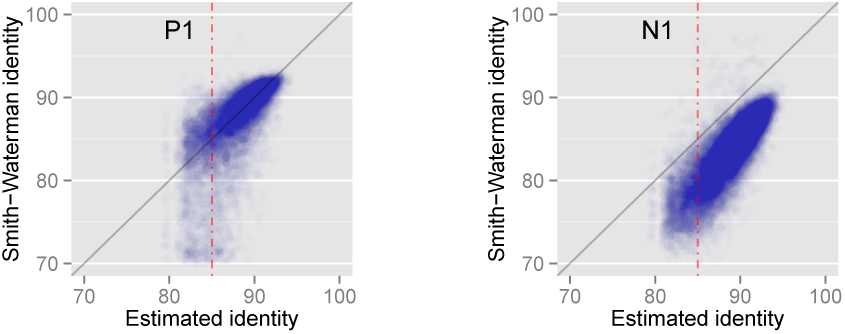
Correlation between Smith-Waterman identity and the identity estimated by mashmap using datasets P1 (PacBio) and N1 (MinION). Red dotted line corresponds to the error cut-off *ϵ_max_* = 0.15.

### 8.3Mapping to RefSeq

We perform mapping of a publicly available PacBio read set consisting of 127,565 reads (each ≥ 5 kbp) sequenced from a mock microbial community containing 20 strains [20]. To demonstrate the scalability of our algorithm, we map these reads against the complete NCBI RefSeq database (838 Gbp) containing sequences from 60,892 organisms. This experiment was executed using default parameters (*l*_0_ = 5000, *ϵ_max_*= 0.15,*p_max_* = 0.001) on an Intel Xeon CPU E7-8837 with 1 TB memory. BWA-MEM and minimap could not index the entire RefSeq database at once with this memory limitation. Mashmap took 29 CPU hours to index the reference and 16 CPU hours for mapping, with a peak memory usage of 660 GB. Note that the same index can be repeatedly used for mapping sequences, conferring our method the ability to process data in real-time. To check the accuracy of our results, we ran BWA-MEM against the 20 known genomes of the mock community. The recall of mashmap against BWA-MEM mappings ranged from 97.7% to 99.1% for all the 20 genomes in the mock community.

## 9 Conclusions

We have presented a fast approximate algorithm for mapping long reads to large reference genomes. Instead of reporting base-level alignments, mashmap reports all reference intervals with sufficient Jaccard similarity compared to the *k*-mer spectrum of the read. In contrast to earlier techniques based on MinHash and winnowing, we provide a formal characterization of the mappings the algorithm is intended to uncover, and provide a provably good algorithm for computing them. In addition, we report an estimate of the alignment error rate tailored to each mapping under an assumed error model. Mashmap provides significant benefits in run-time, memory usage, and scalability, while achieving precision and recall similar to alignment-based methods. Future work aims to extend this method to split-read mapping, compressed reference databases, and additional error models. For example, the *winnowed-minhash* operation could be applied to paths within a de Bruijn graph to recover identity estimates and identify the database sequences most similar to a query sequence. Such approximate algorithms promise to help address the ever increasing scale of genomic data.

## Acknowledgments

This research was supported in part by the Intramural Research Program of the National Human Genome Research Institute, National Institutes of Health, and the U.S. National Science Foundation under IIS-1416259.

